# Shallow nanopore RNA sequencing enables transcriptome profiling for precision cancer medicine

**DOI:** 10.1101/2022.05.31.494109

**Authors:** Andreas Mock, Melissa Braun, Claudia Scholl, Stefan Fröhling, Cihan Erkut

**Affiliations:** Division of Translational Medical Oncology, National Center for Tumor Diseases (NCT) Heidelberg, German Cancer Research Center (DKFZ), Heidelberg, Germany; German Cancer Consortium (DKTK), Heidelberg, Germany; Division of Applied Functional Genomics, DKFZ and NCT Heidelberg, Heidelberg, Germany; Institute of Pathology, Ludwig Maximilians University Munich, Munich, Germany

## Abstract

Transcriptome profiling is a mainstay of translational cancer research and is increasingly finding its way into precision oncology. While bulk RNA sequencing (RNA-seq) is widely available, high costs and long data return time are limiting factors for clinical applications. We investigated a portable nanopore long-read sequencing device (MinION, Oxford Nanopore Technologies) for transcriptome profiling of tumors. In particular, we investigated the impact of lower coverage than that of larger sequencing devices by comparing shallow nanopore RNA-seq data with short-read RNA-seq data generated using reversible dye terminator technology (Illumina) for ten samples representing four cancer types. Coupled with ShaNTi (Shallow Nanopore Sequencing for Transcriptomics), a newly developed data processing pipeline, a turnaround time of five days was achieved. The correlation of normalized gene-level counts between nanopore and Illumina RNA-seq was high for MinION but not for very low-throughput Flongle flow cells (r = 0.89 and r = 0.24, respectively). A cost-saving approach based on multiplexing of four samples per MinION flow cell maintained a high correlation with Illumina data (r = 0.56 – 0.86). In addition, we compared the utility of nanopore and Illumina RNA-seq data for analysis tools commonly applied in translational oncology: (i) Shallow nanopore and Illumina RNA-seq were equally useful for inferring signaling pathway activities with PROGENy. (ii) Highly expressed genes encoding kinases targeted by clinically approved small-molecule inhibitors were reliably identified by shallow nanopore RNA-seq. (iii) In tumor microenvironment composition analysis, quanTIseq performed better than CIBERSORT, likely due to higher average expression of the gene set used for deconvolution. (iv) Shallow nanopore RNA-seq was successfully applied to validate known gene fusions by breakpoint analysis. These findings suggest that shallow nanopore RNA-seq enables rapid, cost-effective, and biologically meaningful transcriptome profiling of tumors and warrants further exploration in precision cancer medicine studies.

## INTRODUCTION

Nanopore sequencing is an emerging third-generation DNA and RNA sequencing (RNA-seq) technology. It is based on the phenomenon that a single DNA or RNA molecule in an electrophysiological solution passes through a nanometer-scale protein pore, accompanied by ions in varying concentrations depending on the nucleotide composition. This causes quantifiable patterns of current fluctuations attributable to the nucleotide sequence [1]. Using the digitized current-level information generated, pretrained artificial neural networks can predict the sequence of very long DNA fragments or full-length transcripts with high accuracy. Because this principle does not require imaging, unlike methods based on reversible dye terminator technology (Illumina), sequencing devices could be substantially downsized [2]. Of particular interest to many investigators is the small, portable MinION sequencer (Oxford Nanopore Technologies), which enables rapid and decentralized sequencing with low investment costs but also has lower throughput than previous methods. In cancer research, MinION-based nanopore sequencing has been successfully employed for mutation detection [3–5], DNA methylome analysis [6–8], DNA copy number profiling [6,9], and the identification of gene fusions [10,11]. In addition, full-length cDNA sequencing has been used to detect aberrant splicing in cancer and to perform differential expression analysis [12,13].

Despite these developments, the full utility of nanopore RNA-seq for transcriptome analysis of human cancers remains elusive. This could be highly relevant for the clinical implementation of precision oncology approaches as the detection of aberrantly expressed genes has the potential to substantially increase the proportion of patients whose care can be individualized based on molecular information compared with DNA-based stratification approaches alone [14]. For example, mRNA expression analysis of the receptor tyrosine kinase genes *FGFR1-3* identified a larger patient population eligible for treatment with the pan-FGFR inhibitor rogaratinib [15]. More broadly, the increasing understanding of the kinome-wide target spectrum of clinically available kinase inhibitors [16] allows the predictive value of kinase expression for response to these drugs to be systematically studied [17–19]. In addition, inferring pathway activities from RNA-seq data is becoming increasingly important as they have been shown to outperform the expression of single genes as biomarkers [20,21]. However, the functional taxonomy of tumor ecosystems based on their transcriptomes is not limited to the cancer cell compartment. For example, deconvolution algorithms are widely applied to estimate the composition of the immune cell microenvironment [22–24].

MinION-based nanopore RNA-seq can be considered “shallow” RNA-seq since its yield is considerably lower than that of standard Illumina RNA-seq. However, predictive *in silico* modeling suggests that biomarkers for precision cancer medicine can be developed with drastically reduced sequencing depth [25]. We investigated the feasibility of using a MinION sequencer for rapid transcriptome profiling of human tumors. In addition to a comparison with matching Illumina RNA-seq data, we explored four applications for precision cancer medicine, i.e., (i) pathway activity inference, (ii) expression of kinases targeted by approved inhibitors, (iii) immune cell deconvolution, and (iv) identification of known fusion genes. Our results show that shallow nanopore RNA-seq enables biologically meaningful transcriptome profiling of tumors and warrants further development as a stratification tool in the clinic.

## METHODS

### Patient samples

Tumor samples from ten patients with rare cancers, i.e., adenoid cystic carcinoma (ACC), dedifferentiated liposarcoma (DDLS), large-cell neuroendocrine carcinoma (LCNC), and synovial sarcoma (SS), were studied. All samples were subjected to quality control, verification of the respective entity, and estimation of tumor cell content by experienced pathologists. RNA was extracted in the Sample Processing Laboratory of the German Cancer Research Center (DKFZ). Illumina RNA-seq had been performed within the MASTER (Molecularly Aided Stratification for Tumor Eradication Research) trial of the National Center for Tumor Diseases (NCT), DKFZ, and the German Cancer Consortium (DKTK) [26]. Library preparation for nanopore RNA-seq was performed with the same analyte previously used for Illumina RNA-seq. Samples were selected based on RNA availability and known entity-defining gene fusions (Table 1).

**Table 1.**
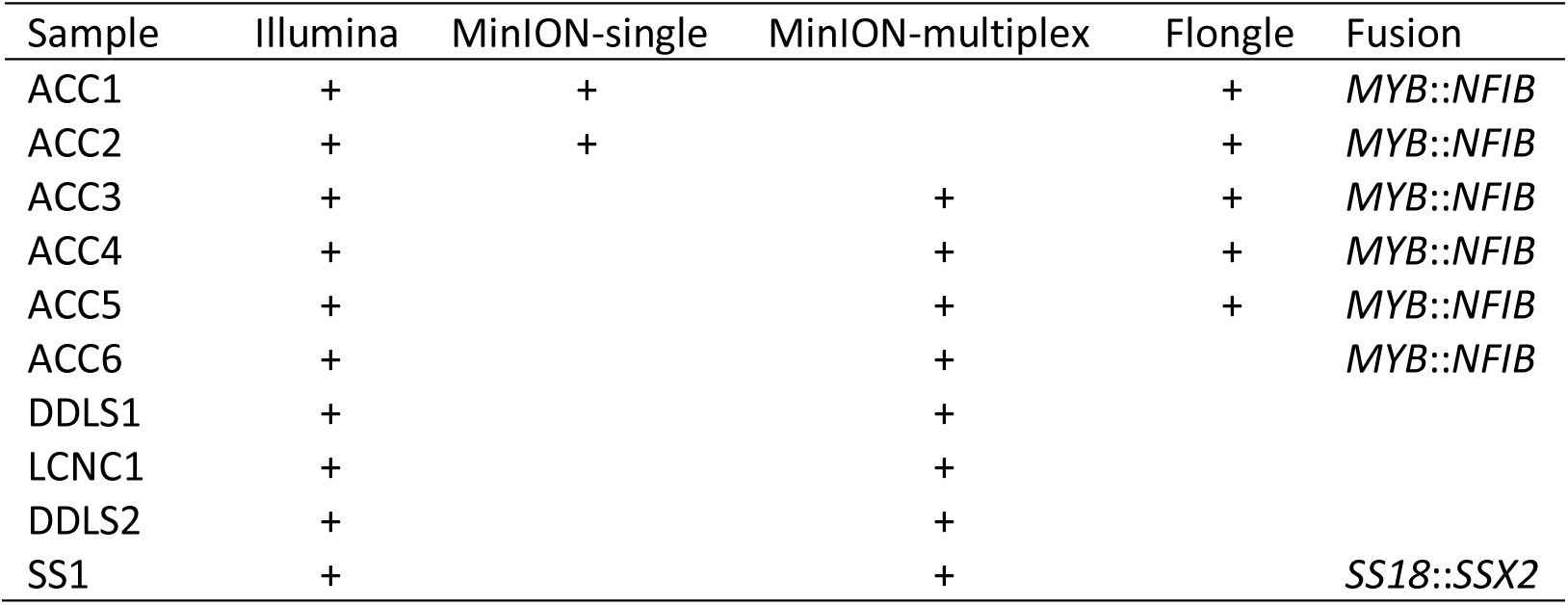
Tumor samples. Listed are the availability of RNA-seq data (indicated by “+”) for the Illumina and various Oxford Nanopore platforms and characteristic gene fusions identified by Illumina-based sequencing.

### Illumina RNA-seq data processing

Reads were processed with the RNA-seq workflow 1.3.0 developed by the DKFZ Omics IT and Data Management Core Facility (https://github.com/DKFZ-ODCF/RNAseqWorkflow). First, FASTQ reads were aligned via two-pass alignment using STAR 2.5.3a [27]. The STAR index was generated from the 1000 Genomes assembly and GENCODE Version 19 gene models with a sjdbOverhang of 200. Alignment call parameters are listed in Table S1. Other parameters were set as default or only pertinent for particular samples. Duplicate marking of the resultant main alignment file was done with sambamba 0.6.5 [28] using eight threads. The chimeric file was sorted using samtools 1.6 [29], and duplicates were marked using sambamba. BAM indexes were also generated using sambamba. Quality control was performed using samtools flagstat [29] and the rnaseqc tool version 1.1.8 [30] with the 1000 Genomes assembly and GENCODE Version 19 gene models. Depth-of-coverage analysis for rnaseqc was turned off. Gene-specific read counting was performed using featureCounts (from Subread 1.5.1) [31] over exon features based on GENCODE Version 19 gene models. Both reads of a paired fragment were used for counting, and the quality threshold was set to 255, indicating that STAR found a unique alignment. Strand-specific counting was also used. For RPKM and TPM calculations, all genes on chromosomes X and Y, the mitochondrial genome, as well as rRNA and tRNA genes were omitted as they are likely to introduce library size estimation biases. All computations were performed on a high-performance compute cluster.

### Nanopore RNA-seq

Direct cDNA sequencing was performed using the SQK-DCS109 kit (Oxford Nanopore Technologies). For analysis of a single sample on a MinION flow cell (version R9.4.1), 5 μg RNA was used as input. For multiplexing on a MinION flow cell, 2.5 μg RNA per sample was used as input, and the native barcoding expansion kit EXP-NBD104 was employed in conjunction with SQK-DCS109. After reverse transcription with Maxima H Minus Reverse Transcriptase (Thermo Scientific), second-strand synthesis was performed using the 2x LongAmp Taq Master Mix (New England Biolabs). The resulting double-stranded cDNA was subjected to end-repair and dA-tailing using the NEBNext Ultra End Repair/dA-Tailing Module (New England Biolabs). For multiplexed libraries, this step was followed by barcode ligation and library pooling. Next, libraries were quantified with a Qubit Fluorometer 3.0 (Life Technologies). Finally, sequencing adapters were added to the library preparations and ligated with Blunt/TA Ligase Master Mix (New England Biolabs), followed by further quality control using a Qubit. Samples ACC1 and ACC2 were analyzed on individual MinION flow cells, while the remaining eight samples were sequenced as multiplexed libraries on two MinION flow cells by pooling four samples for each run. Five ACC samples were also analyzed individually on Flongle flow cells (Table 1). The run time was between 72 and 96 hours, depending on library and flow cell quality.

### Nanopore RNA-seq data processing

We developed a custom pipeline for processing nanopore RNA-seq data, available at https://github.com/cihanerkut/shanti. The workflow includes basecalling, read filtering, demultiplexing (optionally), alignment, and read summarization (Figure S1). Custom parameters for alignment are listed in Table S2. All other parameters were either kept as default or adjusted to the respective sample, e.g., according to the sequencing and barcoding kits used. All computations were performed on a local workstation with 32 threads, 256 GB RAM, and an NVIDIA Tesla V100 16 GB GPU. First, GPU basecalling was performed from FAST5 files with Guppy basecaller 5.0.14 using the super-accuracy model. Adapter trimming was turned off during basecalling of multiplexed data. MinION reads with an average Phred score of less than 7 were filtered out with NanoFilt 2.6.0 [32], as such reads cannot be used for accurate demultiplexing. For Flongle data, a Phred score cutoff of 4 was used due to overall lower basecalling quality. Next, high-quality reads from multiplexed experiments were demultiplexed using Guppy barcoder 5.0.14 applying adapter and barcode trimming. Demultiplexed, filtered, and trimmed reads were aligned to the same 1000 Genomes assembly reference (hs37d5) used for Illumina RNA-seq data preprocessing. Alignment was performed with minimap2 2.22 [33]. Alignment parameters were optimized to achieve the highest rate of reverse-oriented protein-coding genes with the lowest alignment error. SAM-to-BAM conversion, BAM sorting, indexing, and extraction of basic alignment statistics were performed with SAMtools 1.13. Read summarization was performed with featureCounts (from Subread 2.0.3), similar to Illumina data preprocessing. Unstranded and stranded count tables were merged for each sample. Quality reports from basecalled, untrimmed reads before demultiplexing, filtered and demultiplexed reads, and aligned reads were generated using NanoPlot 1.28.2 [32].

### Integration of mRNA abundance data from Illumina and nanopore sequencing

Raw transcript counts from all Illumina and nanopore sequencing runs were combined into one table for downstream analysis, and only the counts of reverse-stranded alignments were used. Illumina sequencing was performed with the TruSeq Stranded mRNA Library Prep system (Illumina), which effectively estimates the abundance of coding cDNA strands. Therefore, only the counts of reverse-stranded alignments were relevant in these data. A similar stranded sequencing approach is not available for nanopore RNA-seq. To estimate the abundance of coding cDNA strands, we used only the counts of reverse-stranded alignments. Similarly, we used the transcripts per million (TPM) values derived from reverse-stranded alignment counts, which normalize raw read counts to the total exon length and library size for each gene and accurately represent mRNA abundance [34]. As a comparable metric for long-read sequencing, we calculated reads per million (RPM) by normalizing reverse-stranded long-read alignment counts to the corresponding library size. Briefly, the total library size was estimated after excluding all genes on chromosomes X and Y, the mitochondrial genome, as well as rRNA and tRNA genes. Gene-wise read counts were divided by the total library size and multiplied by 1,000,000 to calculate RPM values, whereas normalization to total exon length as in TPM calculation was not relevant for long-read sequencing.

### Inference of pathway activities

To infer signaling pathway activities from RNA-seq data, the PROGENy algorithm implemented in the *progeny* R package was used [35]. PROGENy is based on a list of pathway response genes collected from publicly available perturbation experiments. Version 1.10 of the R package contains data on 14 pathways involved in cancer biology (Androgen, EGFR, Estrogen, Hypoxia, JAK-STAT, MAPK, NFkB, p53, PI3K, TGFb, TNFa, Trail, VEGF, and WNT). The input for PROGENy were the TPM and RPM matrices for Illumina and nanopore sequencing, respectively. Pathway scores were scaled to have a mean of 0 and a standard deviation of 1, the default for the function. Different numbers of footprint genes per pathway (default is 100) were investigated.

### Immune cell deconvolution

For immune cell deconvolution from RNA-seq data, CIBERSORT [23] and quanTIseq [24], implemented in the *immunedeconv* R package [36], were employed. CIBERSORT estimates the abundance of 22 immune cell types using 547 genes, whereas in quanTIseq, ten immune cell types are deconvoluted using a set of 170 genes. The input for both algorithms were the TPM and RPM matrices. As recommended by the CIBERSORT authors, quantile normalization was disabled for RNA-seq data, and fractions were calculated in relative mode. QuanTIseq was run with default parameters.

### Visualization of gene fusions

Gene fusions were first confirmed using Integrative Genomics Viewer (IGV) [37]. Subsequently, detailed visualizations with coverage and alignment data were generated using the *Gviz* package in R [38]. Gene models for the GRCh37 human genome assembly were downloaded from ENSEMBL using the BioMart service. Only protein-coding transcripts with the GENCODE Basic tag were used to create a metagene model including all possible exons. Fusion breakpoints were determined by whole-genome sequencing and acquired from the clinical bioinformatics workflow of the MASTER cohort [26].

### Data availability

Sequencing and gene expression data have been deposited at the European Genome-phenome Archive (EGA), which is hosted by the EBI and CRG, under accession number EGAS00001006317.

## RESULTS

### Comparison of shallow nanopore and Illumina RNA-seq for tumor transcriptome profiling

To benchmark shallow nanopore RNA-seq for tumor transcriptome profiling, we analyzed tissue samples from ten patients for whom Illumina RNA-seq data had been generated within the MASTER precision oncology program [26] (Fig. 1a). The workflow from extracted RNA to processed and normalized RNA-seq data took five days for each sequencing run. We automated the standard data preprocessing steps, i.e., basecalling, read filtering, alignment to the reference genome, and read counting, with a custom bioinformatics pipeline that we termed ShaNTi (Shallow Nanopore Transcriptomics) (Fig. S1). In addition to processed data, ShaNTi generates detailed statistics, e.g., regarding sequencing depth, read length, alignment accuracy, etc.

**Fig. 1.**
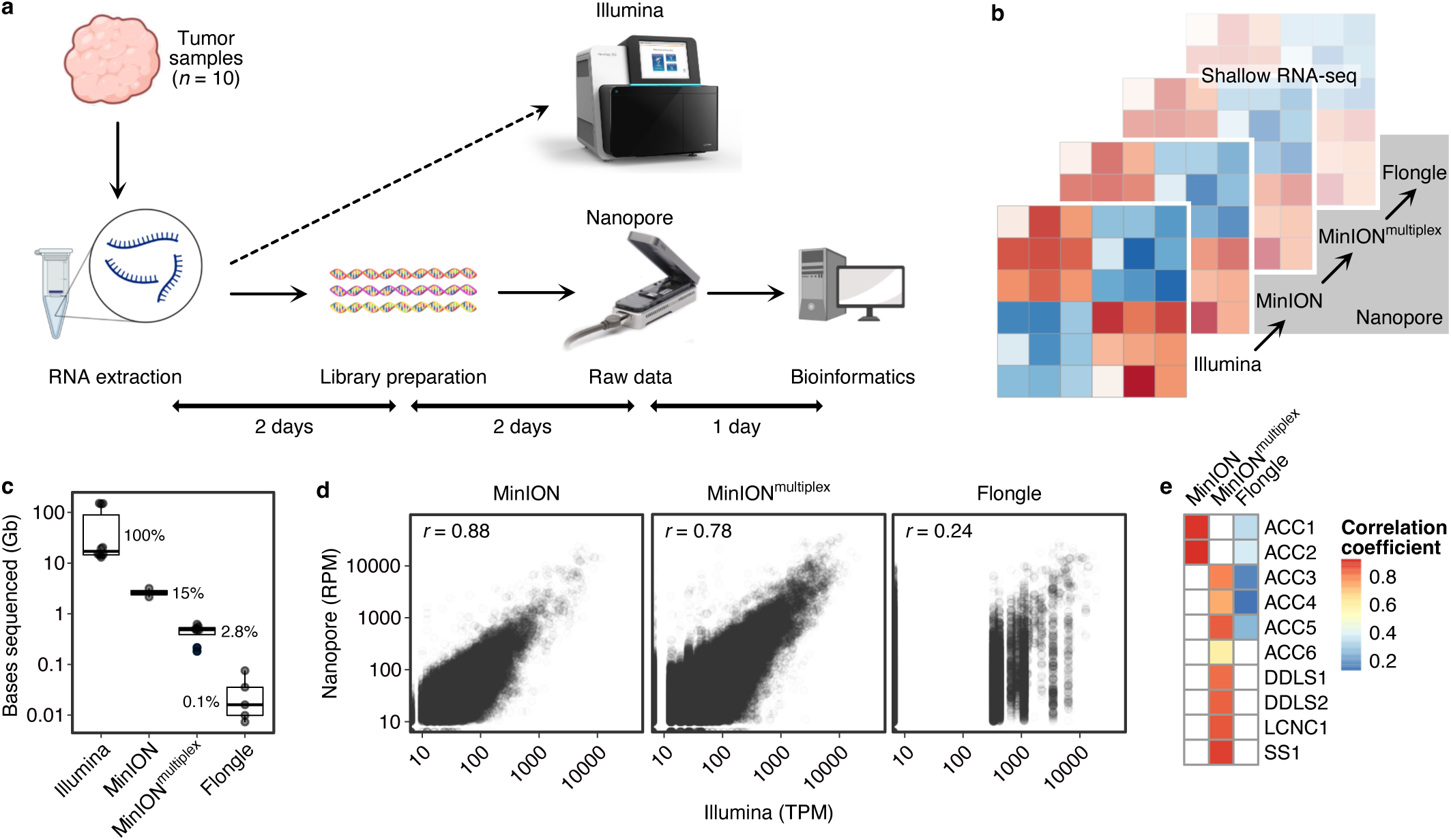
Comparison of shallow nanopore and Illumina RNA-seq. a) Overview of sample and data processing for nanopore RNA-seq (created with Biorender.com). b) Schematic of the different sequencing depths used to perform nanopore RNA-seq (MinION flow cell with one sample, MinION flow cell with four samples [MinION^multiplex^], Flongle flow cell with one sample). c) Comparison of sequencing depth, i.e., bases sequenced, between technologies. Nanopore RNA-seq covered a median of 0.1 – 15% of the bases sequenced with Illumina RNA-seq. d) Correlation of normalized gene-level counts obtained by nanopore (RPM values) and Illumina (TPM values) RNA-seq across all samples and protein-coding genes (Spearman rank correlation coefficient). Each dot represents a gene. e) Correlation of normalized gene-level counts obtained by nanopore and Illumina RNA-seq for individual samples (Spearman rank correlation coefficient).

We examined different sequencing depths (Fig. 1b) and observed that running a single sample on a MinION or Flongle flow cell yielded a median of 15% or 0.1% of the bases sequenced, respectively, compared to Illumina RNA-seq (Fig. 1c). Multiplexing four samples on a MinION flow cell yielded a median of 2.8% of the bases sequenced compared to Illumina RNA-seq (Fig. 1c). The correlation between normalized gene-level counts determined by nanopore or Illumina RNA-seq across samples was high for MinION flow cells (single sample, r = 0.88; four multiplexed samples, r = 0.78) (Fig. 1d) but low when we used a Flongle flow cell (r = 0.24). The correlation between MinION-based nanopore and Illumina RNA-seq was also high for individual samples (single samples, r = 0.88 – 0.89; multiplexed samples r = 0.56 – 0.86) (Fig. 1e). As expected, samples analyzed by nanopore or Illumina RNA-seq formed two distinct clusters in an unsupervised comparison (Fig. S2). Together, these data showed that the transcriptomic measurements obtained by shallow nanopore and Illumina RNA-seq were highly concordant.

### Inference of signaling pathway activities and kinase expression analysis

An emerging application of RNA-seq in precision oncology is the inference of signaling pathway activities from tumor transcriptional profiles, as aberrantly activated signaling cascades provide insight into upstream mutant drivers and associated molecular dependencies that can be exploited therapeutically. To test the use of nanopore RNA-seq data in such an analysis, we applied the PROGENy method [35]. In the original PROGENy framework, 100 so-called footprint genes per pathway were used to compute the activity of a pathway. To simulate the impact of shallow nanopore RNA-seq on the performance of PROGENy, we compared the accuracy of determining an active pathway based on nanopore or Illumina RNA-seq with different coverages. Using the default number of 100 footprint genes per pathway resulted in a drop in performance with decreasing coverage (area under the receiver operating characteristic curve [AUROC], 0.74 – 0.63; Fig. 2a). The same pattern was observed for 200, 300, and 500 footprint genes per pathway. This dependence of PROGENy performance on coverage was no longer present when the number of footprint genes per pathway was set to 1,000 or more (Fig. 2a).

**Fig. 2.**
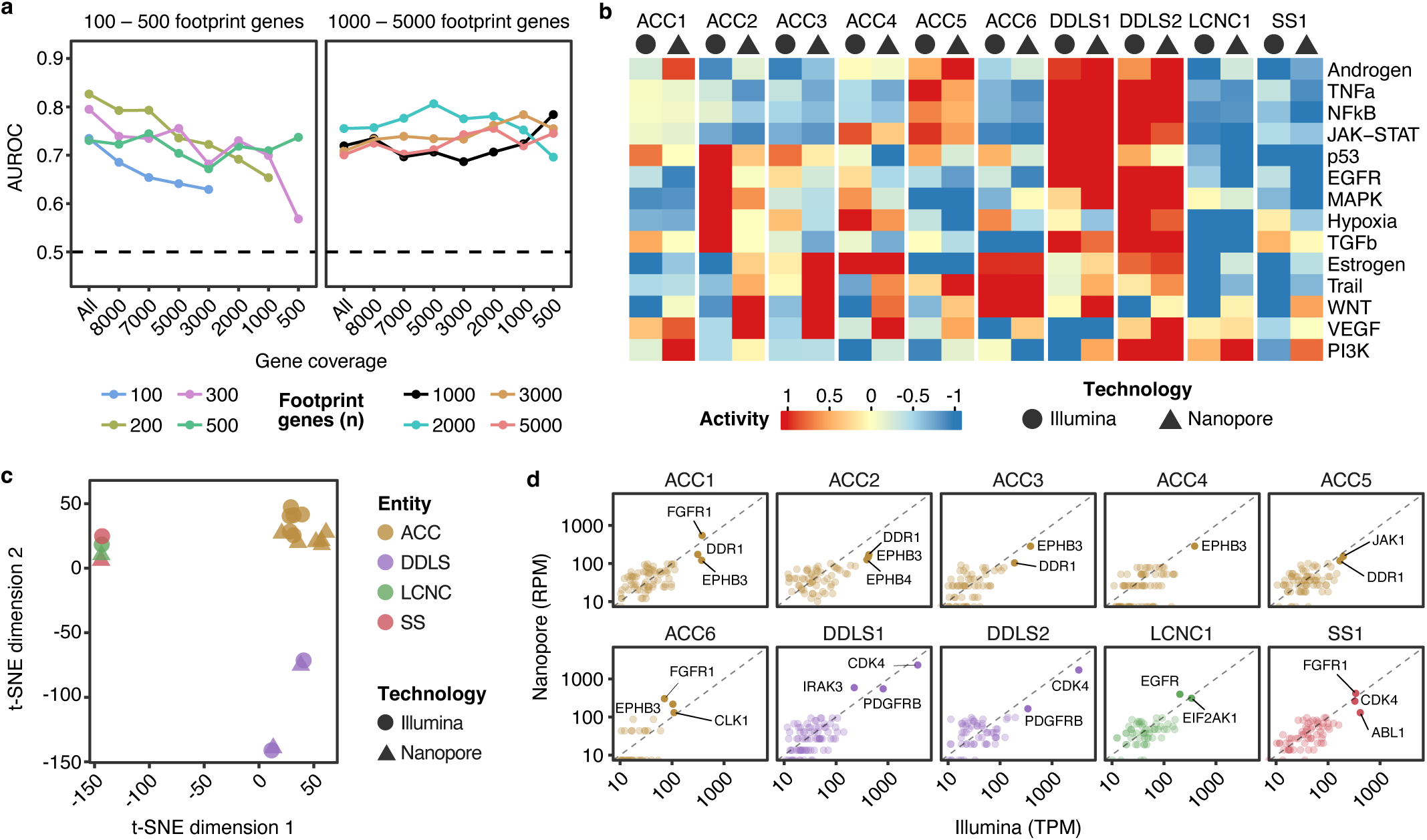
Analysis of pathway activities and kinase expression. a) Performance of PROGENy on nanopore RNA-seq data using different numbers of footprint genes per pathway according to gene coverage. b) Activities of 14 signaling pathways (rows) in ten tumor samples (columns) determined by nanopore (triangle) and Illumina (circle) RNA-seq. c) Visualization of pathway activations shown in panel b after dimensionality reduction with t-distributed stochastic neighbor embedding (t-SNE). d) Correlation of expression of kinases in the target spectrum of clinically approved inhibitors determined by nanopore and Illumina RNA-seq. For each sample, the most highly expressed kinases are labeled.

Running PROGENy with these optimized parameters, i.e., 1,000 footprint genes, resulted in an agreement between pathway activity inferences based on nanopore and Illumina RNA-seq data (AUROC, 0.72; Fig. 2b). The two DDLS samples showed activity of more pathways than the ACC, LCNC, and SS samples. The results were consistent with published results on recurrent mutations or aberrant gene expression driving activation of specific pathways in the respective entities, e.g., “PI3K” in LCNC [39,40], “Androgen” in ACC [41,42], and “NFkB” in DDLS [43]. Unsupervised dimensionality reduction of pathway activation scores showed high similarity of ACC transcriptomes with minor differences attributable to sequencing technology (Fig. 2c), recapitulating the concordance of pathway activity inferences based on nanopore and Illumina RNA-seq data. The other entities clustered separately but independent of sequencing technology, indicating that ACC, DDLS, LCNC, and SS are characterized by distinct transcriptional networks.

In addition to studying entire signaling pathways, we also investigated whether nanopore RNA-seq can accurately detect highly expressed individual kinase genes, which could be potential oncogenic drivers and exploited for kinase-targeted therapies. We, therefore, performed comparative expression analyses of 144 kinase genes that fall within the target spectrum of the 33 clinical kinase inhibitors approved by the United States Food and Drug Administration [16]. We found that in all samples, the kinase genes with the highest expression levels could be reliably identified by nanopore RNA-seq (Fig. 2d). In ACC samples, *FGFR1, EPHB3*, and *DDR1* were recurrently expressed at high levels (Fig. 2d). FGFR family members are commonly upregulated in ACC, and treatment with FGFR inhibitors has shown clinical efficacy in this disease [44–46]. The two DDLS samples were characterized by high *CDK4* and *PDGFRB* expression (Fig. 2d). *CDK4* is amplified in more than 90% of DDLS cases, and clinical trials have shown that the CDK4 inhibitors palbociclib and abemaciclib had a favorable effect on progression-free survival [47,48]. Together, these results demonstrate that nanopore RNA-seq can be used to reliably determine signaling pathway activity and kinase gene expression.

### Immune cell deconvolution

Understanding the tumor microenvironment is becoming increasingly important in precision cancer medicine. For example, the prediction of response to immune checkpoint inhibitors is aided by knowledge of immune cell fractions, whose abundance can be estimated by deconvolution algorithms from bulk RNA-seq data. We employed two of the most widely used algorithms, CIBERSORT [23] and quanTIseq [24]. In CIBERSORT, deconvolution is based on 547 genes covering the profiles of 22 immune cell types. In comparison, quanTIseq uses 170 genes to detect ten immune cell types. To assess the applicability of the two methods to shallow RNA-seq data, we first compared the average expression of CIBERSORT and quanTIseq gene sets in the Illumina and nanopore RNA-seq data (Fig. 3a). In both Illumina and nanopore RNA-seq data, the CIBERSORT reference gene set was significantly lower expressed compared to all protein-coding genes (p < 0.001), whereas no difference was observed for the quanTIseq gene set (p > 0.05). Accordingly, the fraction of genes with zero counts in more than 50% of samples was higher in the CIBERSORT than in the quanTIseq gene set (Fig. 3b; Illumina, 5.5% vs. 1.8%; nanopore, 64.7% vs. 44.8%). In contrast to Illumina RNA-seq, shallow nanopore RNA-seq cannot detect transcripts of very low expressed genes. Using such genes for immune cell deconvolution based on shallow RNA-seq data may therefore lead to incorrect estimates. Indeed, the average abundance values based on Illumina and shallow nanopore RNA-seq data were considerably more similar when using quanTIseq instead of CIBERSORT (Fig. 3c,d, Fig. S3).

**Fig. 3.**
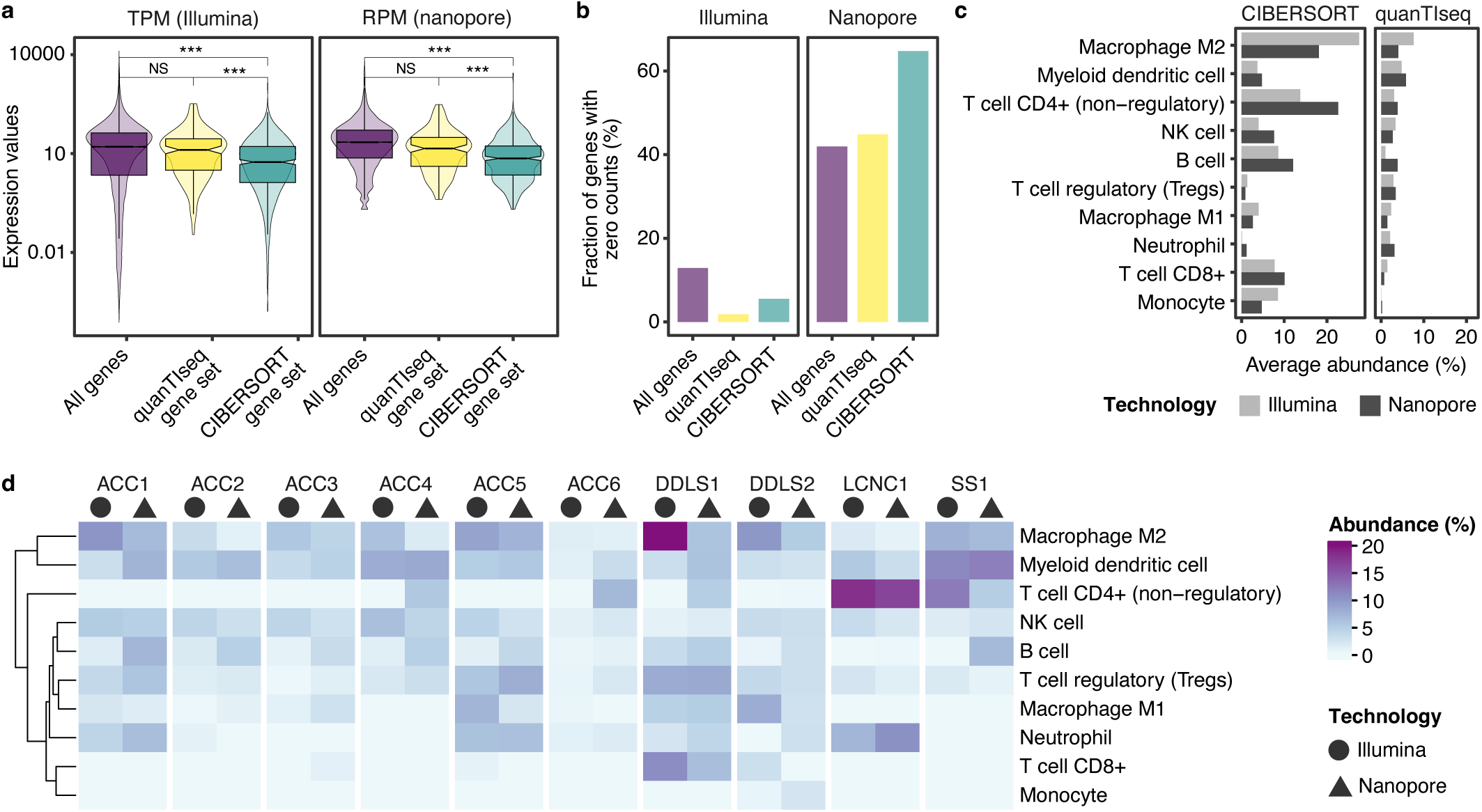
Estimation of immune cell abundance from shallow nanopore and Illumina RNA-seq data. a) Expression, determined by Illumina and nanopore RNA-seq, of all protein-coding genes and gene sets used in quanTIseq (n = 170) and CIBERSORT (n = 547). Groups were compared using Welch’s t-test. ***, p < 0.001; NS, not significant. b) Number of genes with zero counts in more than 50% of samples, as determined by Illumina and nanopore RNA-seq, of all protein-coding genes and gene sets used in quanTIseq (n = 170) and CIBERSORT (n = 547). c) Estimated average abundance of immune cell fractions based on Illumina and shallow nanopore RNA-seq data as a function of the deconvolution method used. d) Immune cell abundance of the ten tumor samples calculated with quanTIseq based on Illumina and nanopore RNA-seq data.

We, therefore, used quanTIseq for subsequent analyses. In the six ACC cases, both sequencing technologies identified immune-excluded microenvironments, as indicated by the low abundance of all immune cell types, consistent with a recent study [49] (Fig. 3d). The LCNC case displayed a high frequency of CD4+ T cells, again based on both Illumina and shallow nanopore RNA-seq data, which has been associated with unfavorable recurrence-free survival in LCNC of the lung [50]. In conclusion, quanTIseq can most likely be used to deconvolute immune cell fractions from shallow nanopore RNA-seq data; however, larger sample size is required to confirm this assumption.

### Visualization of gene fusions

Many cancers are associated with recurrent gene fusions that drive tumorigenesis and may also be diagnostic biomarkers, such as *SS18*::*SSX* in synovial sarcoma [51]. Predicting and visualizing gene fusions based on short-read transcriptome data is challenging because split reads spanning fusion breakpoints carry limited information about the corresponding genomic coordinates. Since the information content of long reads is much higher in this regard, technologies such as nanopore RNA-seq are better suited for these tasks [52]. Although the current release of our ShaNTi bioinformatics pipeline cannot yet automatically detect gene fusions, we aimed to identify them manually using primary, i.e., longest and most accurate, alignments generated by the pipeline and by interactively visualizing the genomic positions of the corresponding complementary alignments in IGV. To test this strategy, we used the ACC samples, which all harbor a disease-defining *MYB*::*NFIB* fusion consisting mainly of portions of the gene encoding the MYB transcription factor [53–55]. With this prior knowledge, we visualized the primary alignments with *MYB* and *NFIB* of the long and short reads obtained from the analysis of ACC samples, supported by knowledge of the exact genomic breakpoints in these genes obtained from whole-genome sequencing of this sample in the MASTER program (Fig. S4-S9).

In a representative sample (ACC2), we examined the 5’ portion of the fusion consisting of portions of the *MYB* gene and observed that both short and long *MYB* reads terminated prematurely near the breakpoint (Fig. 4). The coverage of all exons upstream of the breakpoint was high for both sequencing methods, confirming the expression of this region of *MYB*. In addition, there were reads, albeit few, that aligned to the exons downstream of the breakpoint, which eliminated the possibility that *MYB* is affected by a large deletion. Thus, the primary alignments indicated the presence of a chimeric transcript that involved *MYB*.

**Fig. 4.**
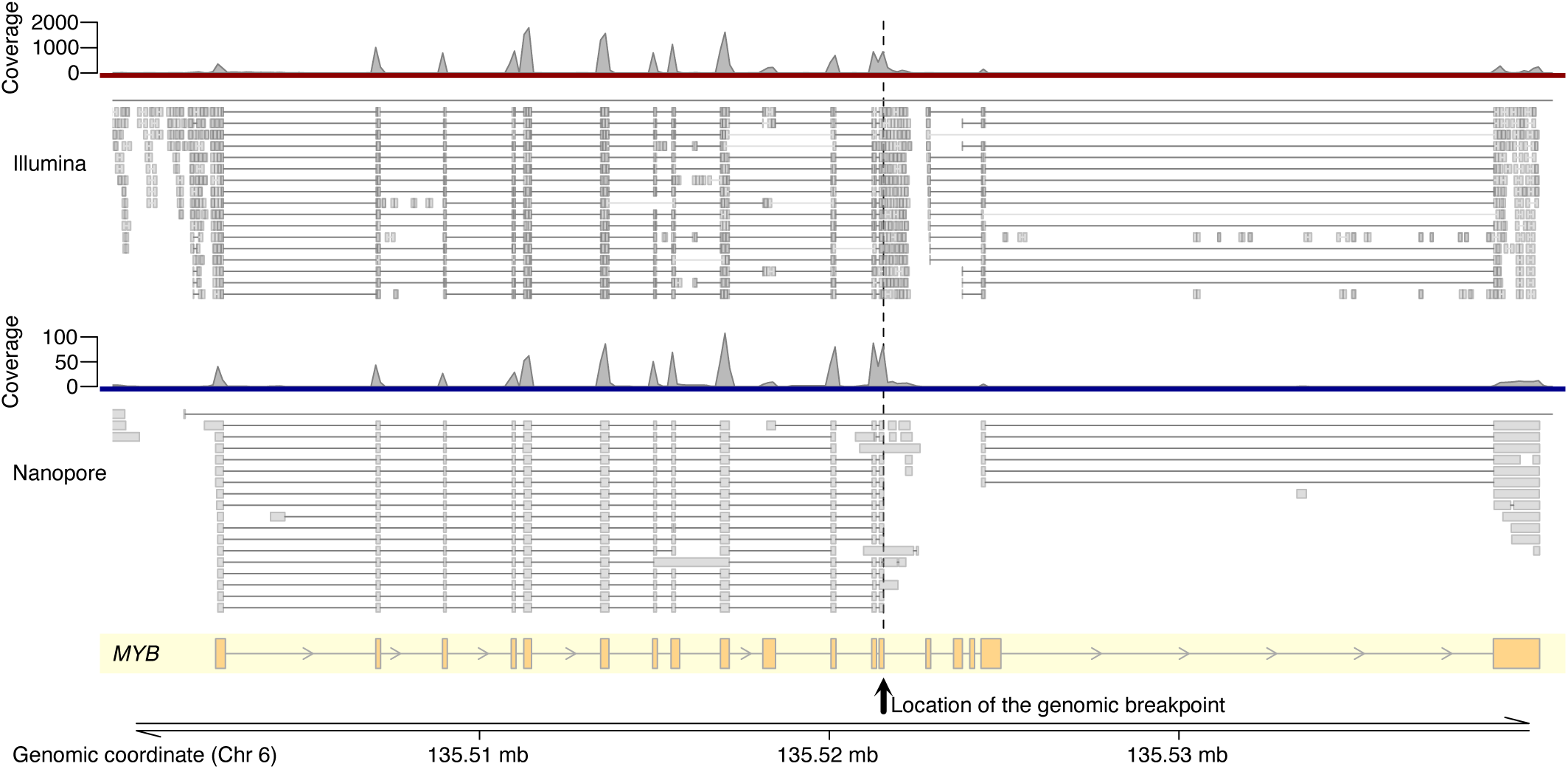
Visualization of short- and long-read alignments to detect gene fusions. Short (top) and long (middle) reads obtained from Illumina and shallow nanopore RNA-seq, respectively, of sample ACC2, plotted as alignments to the *MYB* locus (bottom). For each read, aligned regions are represented by filled boxes, whereas spliced introns are shown as thin lines. The discontinuity of alignments downstream of the breakpoint indicates the end of the *MYB* portion of the chimeric transcript. The corresponding quantitative pileup results representing read coverage are plotted as bar graphs above alignments. The black arrow and the dashed line indicate the position of the genomic breakpoint detected by whole-genome sequencing. In the representation of the gene model (bottom), arrowheads, boxes, and lines indicate the direction of transcription, exons, and introns, respectively.

The 3’ portion of *MYB*::*NFIB* contains only a short coding sequence of *NFIB*, making it more difficult to visualize the *NFIB* breakpoint from primary alignments (Fig. S4). Focusing on nanopore data, we used the supplementary alignments to *MYB* primary alignments and observed that about 70% of them mapped to the *NFIB* gene. Unlike short reads, long reads approximate full-length transcripts. Therefore, a small number of supplementary alignments of long split reads can already be very informative about a genomic fusion event. The fact that most of these supplementary alignments of *MYB* were localized to *NFIB* indicated that chimeric transcripts of these two genes exist in the transcriptome.

However, it must be kept in mind that this approach is sensitive to gene expression levels. When we applied it to the synovial sarcoma sample (SS1) harboring an *SSX2::SS18* fusion gene, we observed few split reads because *SSX2* expression in the sample was very low (Fig. S10). Together, these results indicate that the shallow RNA-seq data generated by our workflow can be used to identify gene fusions, with confidence dependent on the number of covering long reads.

## DISCUSSION

The low cost of setting up and running a MinION sequencer “democratizes” the use of long-read sequencing in academic laboratories. To assess the performance of this new technology in translational oncology, we generated shallow nanopore RNA-seq data using tumor RNA previously subjected to Illumina RNA-seq within a precision oncology program. Our comparative analysis of long-read nanopore and short-read Illumina RNA-seq data demonstrates the feasibility of implementing shallow RNA-seq using the MinION sequencer for transcriptome profiling of human cancers. To address the challenge of a dedicated, easy-to-use bioinformatics workflow for this application, we developed ShaNTi, an automated pipeline that provides a turnaround time from extracted RNA to processed and normalized expression data of only five days. The data preprocessing workflow starts with raw current-level data and creates alignments, transcript quantifications, and quality controls automated. ShaNTi is publicly available and uses exclusively open-source software and can thus be easily adapted to any workstation equipped with the necessary hardware.

To determine the optimum sequencing depth required to achieve a high correlation with Illumina RNA-seq data, we considered different flow cells and multiplexing of samples. Specifically, the comparison was made for a single sample per MinION flow cell, multiplexing four samples on a MinION flow cell, and a single sample on the smaller and less expensive Flongle flow cell. The best compromise between sequencing depth and cost was reached when multiplexing four samples on a MinION flow cell. This result is consistent with a comprehensive computational simulation of RNA-seq data generated within The Cancer Genome Atlas initiative, which showed that a 10-to 100-fold reduction in sequencing depth resulted in no change in the performance of predicting patient outcomes based on transcriptome profiling [25].

Following this technical benchmarking, we pursued a first translational application and derived signaling pathway activities from nanopore RNA-seq data and compared them with the corresponding Illumina RNA-seq results. To this end, we applied PROGENy, an emerging methodology used to better understand oncogenic signaling and, of particular relevance to precision oncology, to relate pathway activities to drug responses [35,56,57]. Based on single-cell RNA-seq data, it has been shown that the number of footprint genes used to infer the activity of a given pathway needs to be tuned [58]. We found that such parameter tuning is also crucial for our shallow RNA-seq approach, as the performance of PROGENy dropped with decreasing gene coverage when we included a smaller number of footprint genes per pathway (100-500), while it remained stable when we included 1,000 genes or more. Using this optimized parameter, we observed high concordance between the results derived from nanopore and Illumina RNA-seq, consistent with previous data on the activity of specific pathways in the entities studied, e.g., androgen signaling in ACC [41,42]. However, because our study is the first application of PROGENy in ACC, LCNC, DDLS, and SS, a more comprehensive comparison is not possible.

In addition to pathway activity, we determined in each sample the most highly expressed kinase genes whose protein products are within the target range of clinically approved inhibitors and observed that findings based on Illumina RNA-seq were confirmed in all cases by nanopore RNA-seq. For example, FGFR family members were commonly overexpressed in ACC, consistent with the clinical efficacy of FGFR blockade in this entity [44–46]. Similarly, the DDLS sample showed *CDK4* overexpression, which, due to amplification of the *CDK4* locus, occurs in more than 90% of DDLS cases and provides a rationale for using CDK4 inhibitors, which have been associated with prolonged progression-free survival in clinical trials [47,48].

Next, we explored the application of nanopore RNA-seq to predict the tumor microenvironment composition using the CIBERSORT and quanTIseq algorithms. Because the shallow RNA-seq approach cannot capture sparsely-expressed genes, we first investigated the average expression of the gene sets used in CIBERSORT and quanTIseq in our Illumina RNA-seq data and observed that the reference genes used in CIBERSORT were significantly less expressed and more often undetectable than those used in quanTIseq. This is a potential reason why the concordance between the immune cell estimates derived from Illumina and nanopore RNA-seq data was inferior when using CIBERSORT. In line with a recent study [49], the six ACC cases displayed an immune-excluded microenvironment, with an overall low fraction of immune cells. The high proportion of CD4+ T cells in the LCNC case may be an unfavorable prognostic factor for recurrence-free survival, as previously described [50]. Further studies correlating immune cell fractions estimated by quanTIseq with immunohistochemical staining of the same tumor tissue are needed to validate these findings.

In a third application of diagnostic and therapeutic relevance, we demonstrated that shallow nanopore RNA-seq data can be used to detect and visualize gene fusions using read alignments. In the interest of rapid turnaround, which can be critical in certain clinical situations, automatic detection of gene fusions is desirable, requiring further development of our pipeline for processing long-read sequencing data.

In summary, we showed that shallow nanopore RNA-seq enables biologically meaningful transcriptome profiling of human cancers and thus has the potential to complement short-read-based sequencing workflows, especially in applications where rapid processing is required. Our next steps will be to develop our workflow further and, in particular, to test it prospectively alongside the diagnostic workup of the MASTER precision oncology trial.

## ACKNOWLEDGEMENTS

We thank Stefanie Reinhart, Tatjana Walther, and Lena Figur for technical assistance; Roman Doll for his contribution to the development of the ShaNTi pipeline; the NCT/DKFZ Sample Processing Lab for providing RNA samples; the DKFZ Omics IT and Data Management Facility for clinical bioinformatics workflows; and the NCT/DKFZ/DKTK MASTER team and all members of Scholl and Fröhling groups for valuable discussions. This project was supported by endowment funds (Stiftung für Krebs-und Scharlachforschung) of the Medical Faculty of Heidelberg University. A.M. was the recipient of a fellowship of the Physician-Scientist Program of the Medical Faculty of Heidelberg University.

## AUTHOR CONTRIBUTIONS

A.M. and C.E. conceptualized the study and wrote the paper. A.M., M.B., and C.E. performed nanopore sequencing and collected and analyzed the data. C.E. created the bioinformatics pipeline. A.M., C.S., S.F., and C.E. interpreted the data. All authors reviewed and commented on the paper.

