## Supplemental data for "Shallow nanopore RNA sequencing enables transcriptome profiling for precision cancer medicine"

German Cancer Research Center

Im Neuenheimer Feld 581

69120 Heidelberg

### SUPPLEMENTARY INFORMATION

#### Table S1. STAR alignment parameters

| Parameter | Value |
|---|---|
| --alignIntronMax | 1100000 |
| --alignIntronMin | 20 |
| --alignMatesGapMax | 1100000 |
| --alignSJstitchMismatchNMax | 5 -1 5 5 |
| --alignSJDBoverhangMin | 3 |
| --chimJunctionOverhangMin | 15 |
| --chimScoreMin | 1 |
| --chimScoreJunctionNonGTAG | 0 |
| --chimSegmentMin | 15 |
| --chimSegmentReadGapMax | 3 |
| --clip3pAdapterSeq | AGATCGGAAGAGCACACGTCTGAACTCCAGTCA |
| --genomeLoad | NoSharedMemory |
| --limitBAMsortRAM | 100000000000 |
| --outBAMsortingThreadN | 1 |
| --outSAMstrandField | intronMotif |
| --outSAMtype | BAM Unsorted SortedByCoordinate |
| --outSAMunmapped | Within KeepPairs |
| --outFilterMismatchNmax | 5 |
| --outFilterMismatchNoverLmax | 0.3 |
| --outFilterMultimapNmax | 1 |
| --readFilesCommand | gunzip -c |
| --runThreadN | 8 |
| --sjdbOverhang | 200 |
| --twopass1readsN | -1 |
| --twopassMode | Basic |

#### Table S2. Minimap2 alignment parameters

| Parameter | Value |
|---|---|
| -x | splice |
| -2 | |
| -a | |
| -k | 28 |
| -I | 100G |
| -w | 30 |
| -t | 32 |
| -K | 1G |
| -L | |
| --cs | long |
| --sam-hit-only | |
| --MD | |

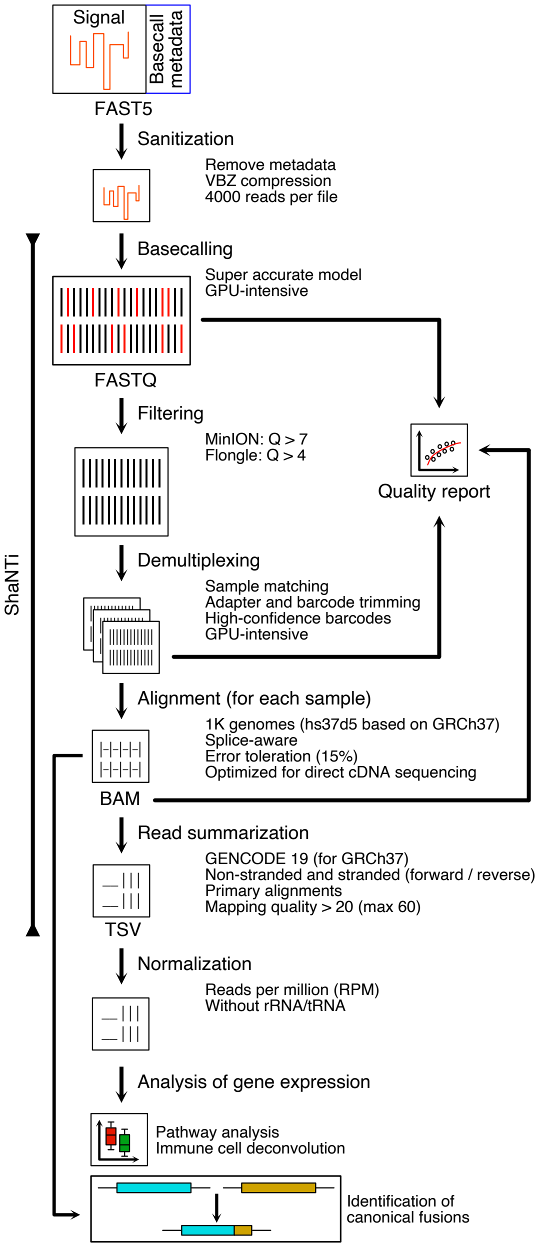


**Fig. S1.** Data processing workflow for tumor transcriptome profiling by shallow nanopore RNA-seq. Current-level data are recorded in FAST5 files during sequencing. Metadata not needed for downstream analysis are removed, and signal data are compressed with the VBZ algorithm to generate standardized FAST5 files, which are used for basecalling with the "super accurate" model developed by Oxford Nanopore Technologies. The resulting sequence data are saved into FASTQ files. Low-quality reads based on the Phred score (indicated by red bars) are filtered out before further processing. For barcoded experiments, filtered FASTQ files are demultiplexed into individual samples and saved as separate FASTQ files. For each sample, reads are aligned to a reference genome, and the output is saved into sorted and indexed BAM files, which are used for read summarization, i.e., quantification of gene expression. The resulting counts are normalized to library size to yield calculated RPM values, which are used to analyze gene expression. In parallel, BAM files after alignment can be used to identify canonical gene fusions employing a visualization software such as IGV.

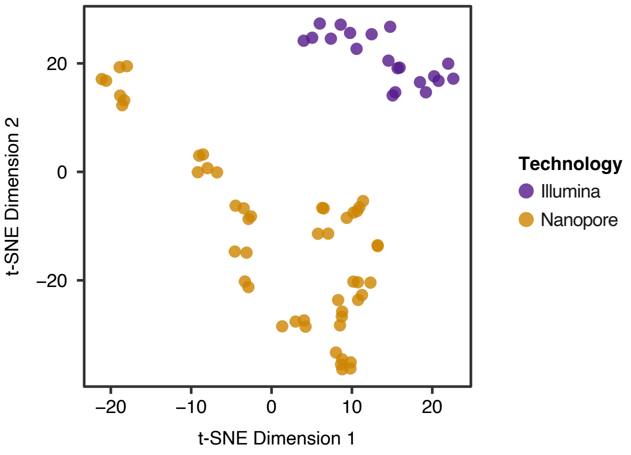


**Fig. S2.** Visualization of TPM (Illumina RNA-seq) and RPM (shallow nanopore RNA-seq) values of all protein-coding genes using t-SNE.

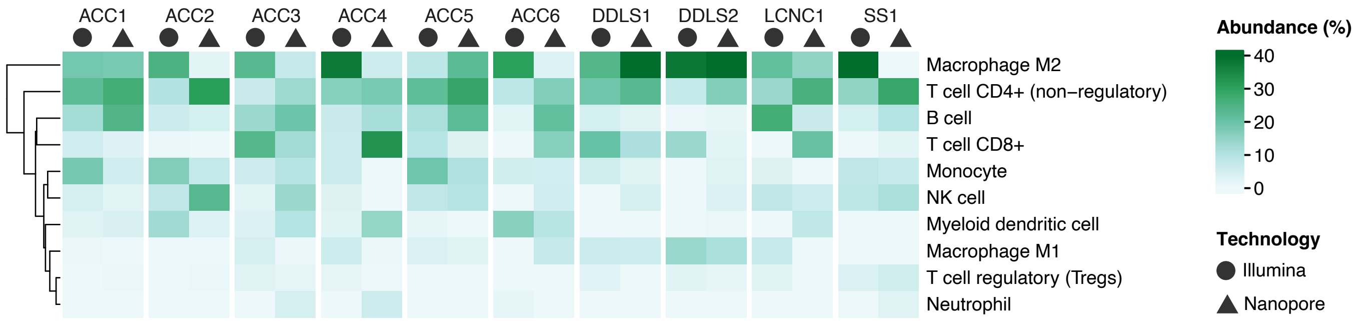


**Fig. S3.** Immune cell abundance of ten tumor samples calculated with CIBERSORT based on Illumina and nanopore RNA-seq data.

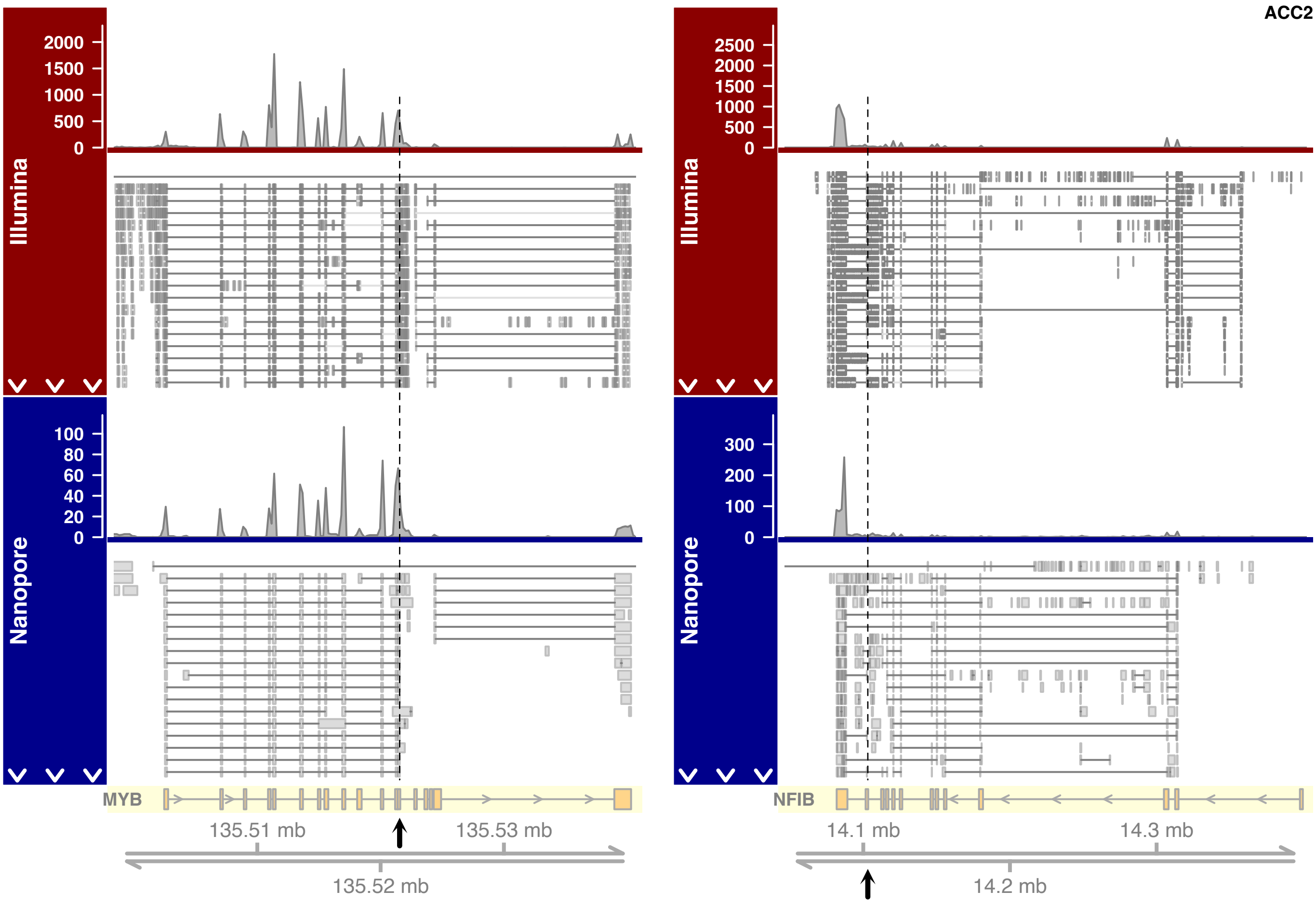


**Fig. S4.** Visualization of short- and long-read alignments to *MYB* and *NFIB* in patient sample ACC2. Short (top) and long (middle) reads obtained from Illumina and shallow nanopore RNA-seq, respectively, of sample ACC2, plotted as alignments to the *MYB* (left) and *NFIB* (right) loci (bottom). For each read, aligned regions are represented by filled boxes, whereas spliced introns are shown as thin lines. The corresponding quantitative pileup results representing read coverage are plotted as bar graphs above the alignments. Black arrows indicate the position of the genomic breakpoint detected by whole-genome sequencing. In the representation of the gene model (bottom), arrowheads, boxes, and lines indicate the direction of transcription, exons, and introns, respectively.

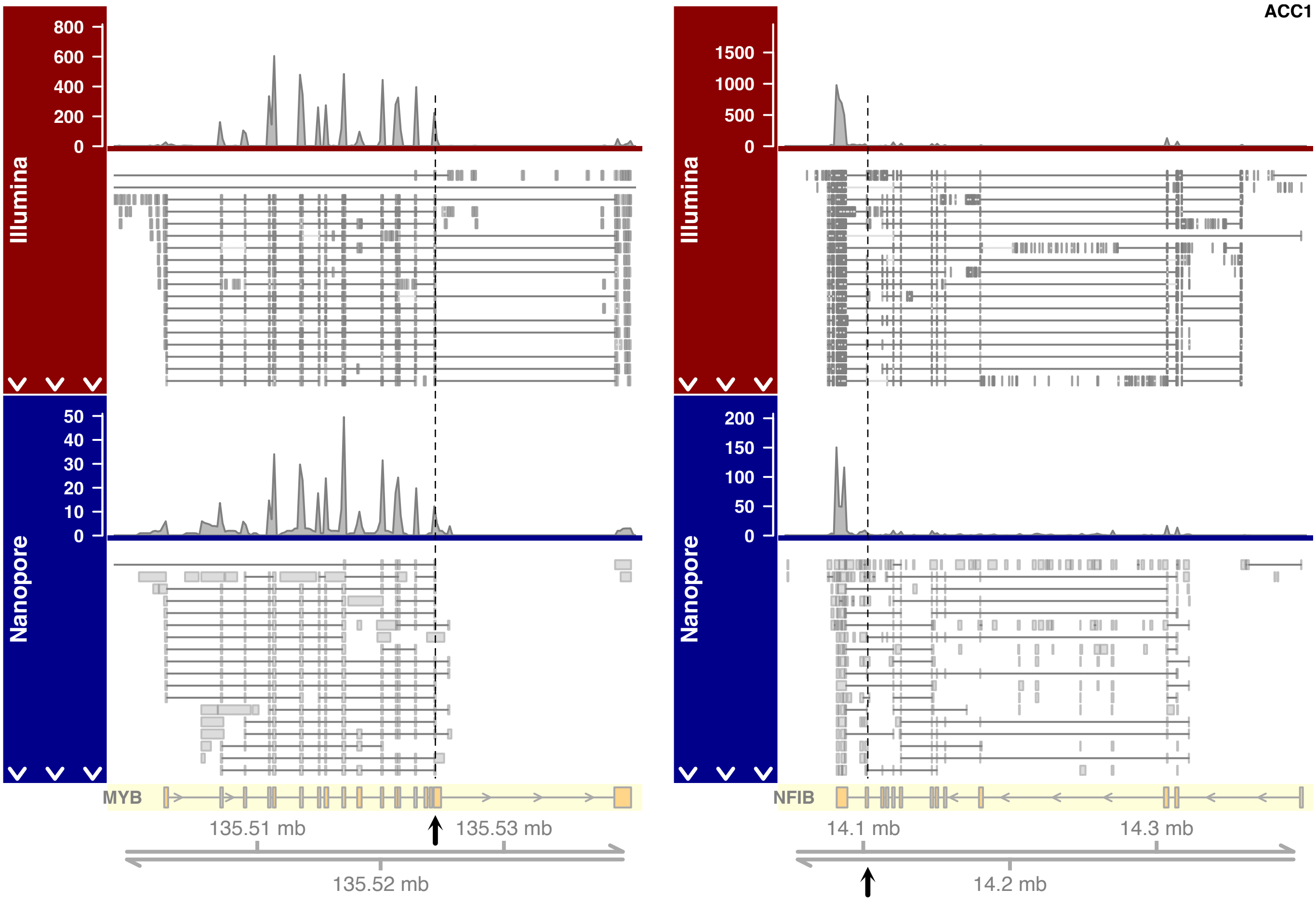


**Fig. S5.** Visualization of short- and long-read alignments to *MYB* and *NFIB* in patient sample ACC1. See Fig. S4 for description.

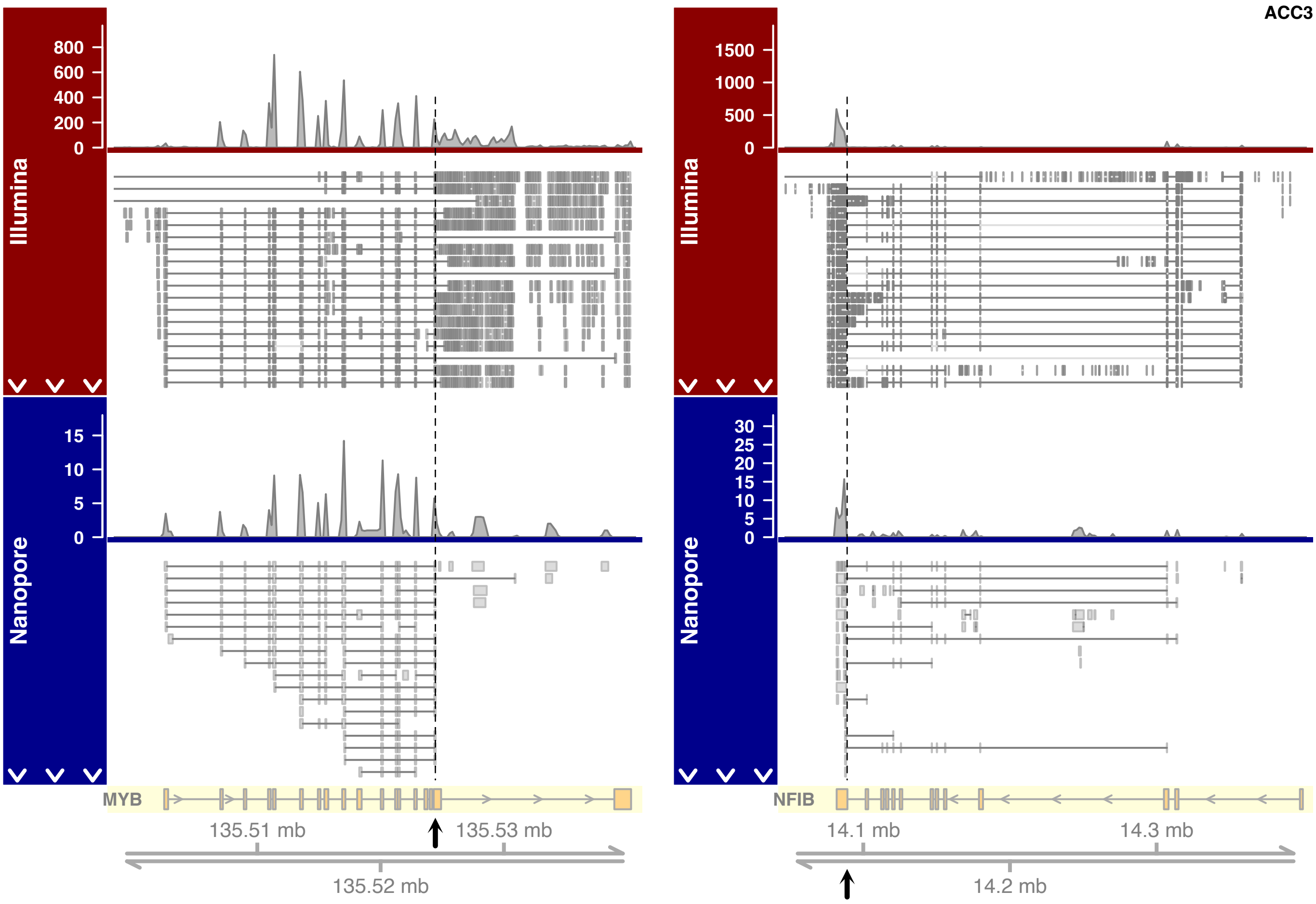


**Fig. S6.** Visualization of short- and long-read alignments to *MYB* and *NFIB* in patient sample ACC3. See Fig. S4 for description.

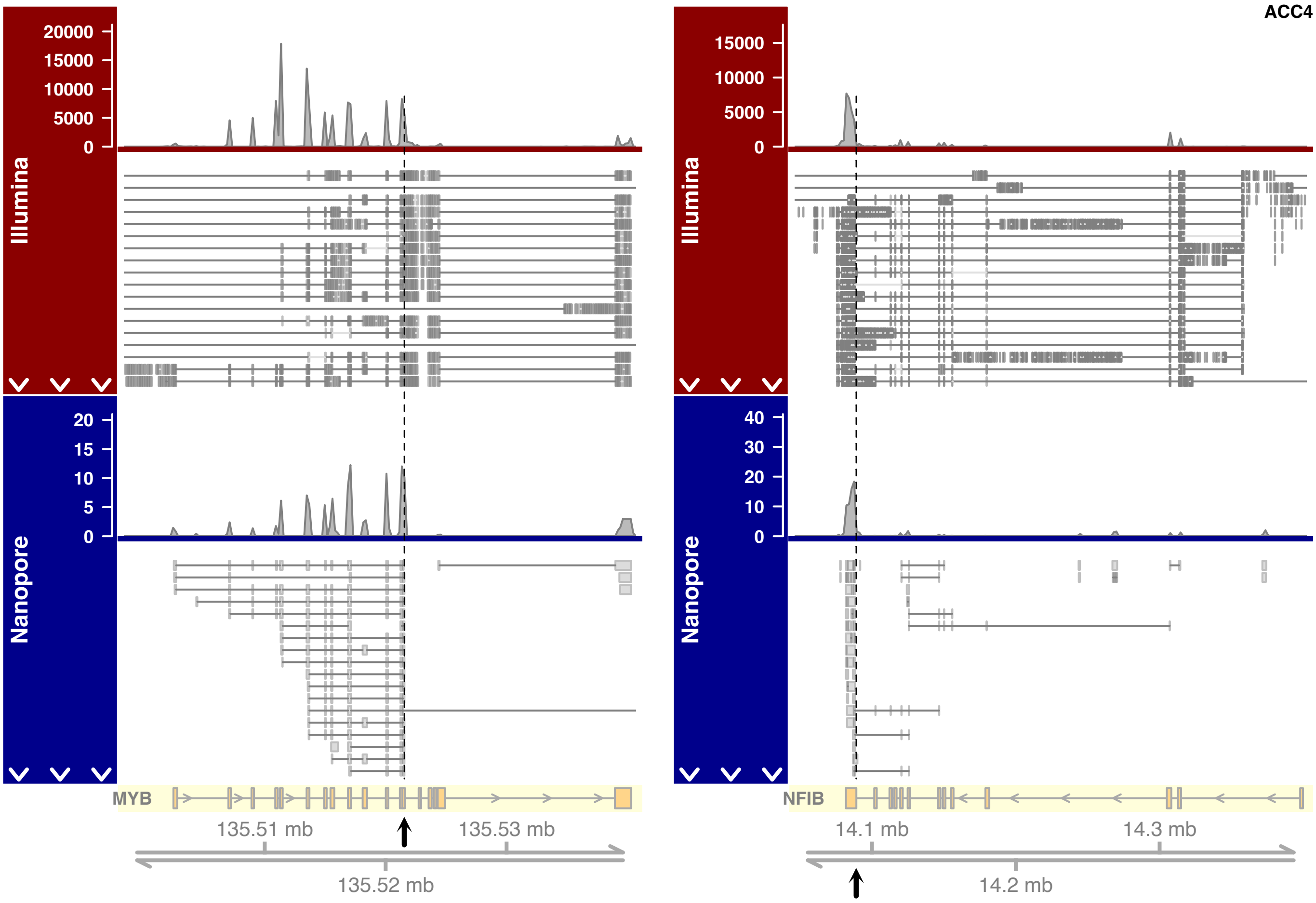


**Fig. S7.** Visualization of short- and long-read alignments to *MYB* and *NFIB* in patient sample ACC4. See Fig. S4 for description.

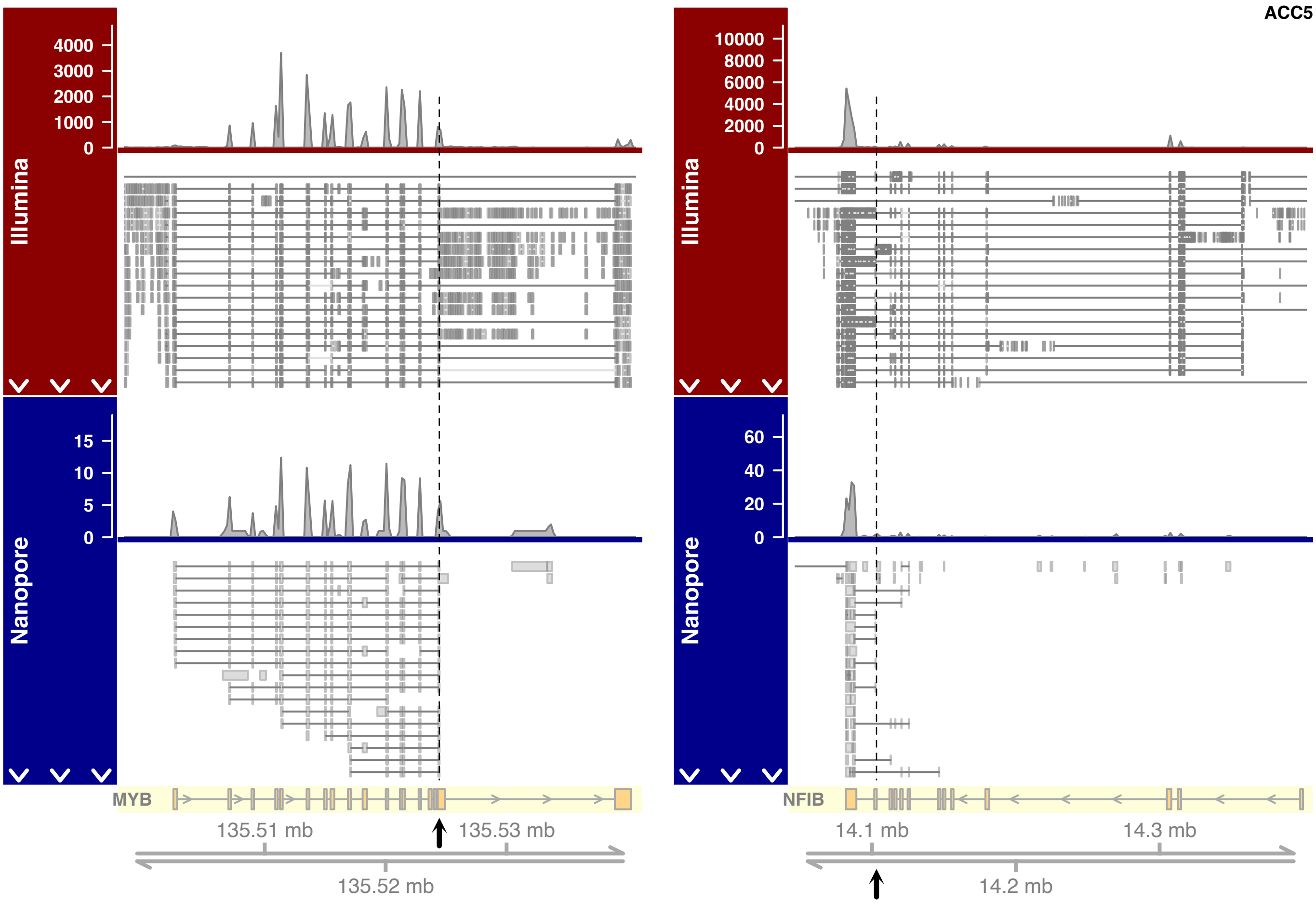


**Fig. S8.** Visualization of short- and long-read alignments to *MYB* and *NFIB* in patient sample ACC5. See Fig. S4 for description.

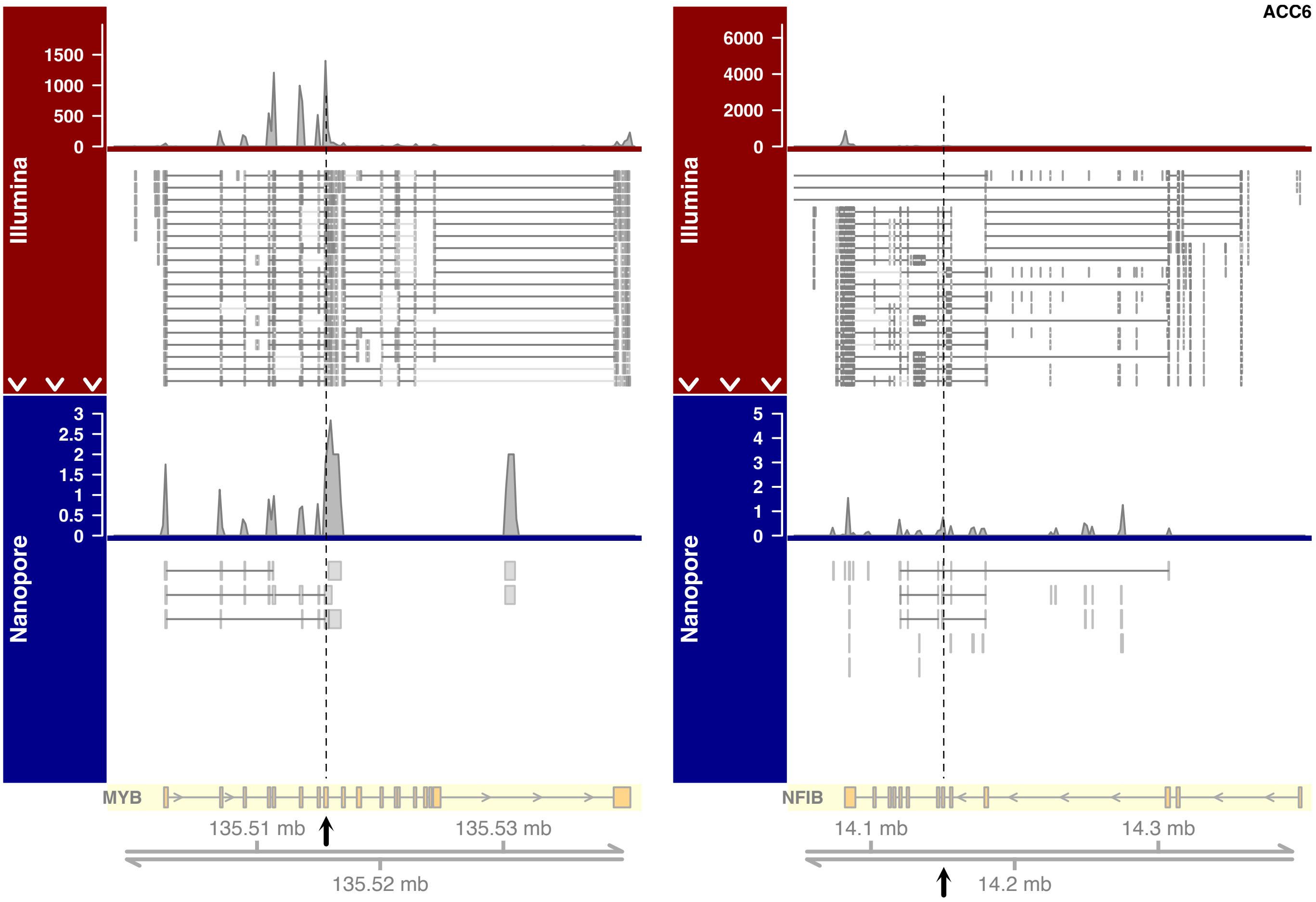


**Fig. S9.** Visualization of short- and long-read alignments to *MYB* and *NFIB* in patient sample ACC6. See Fig. S4 for description.

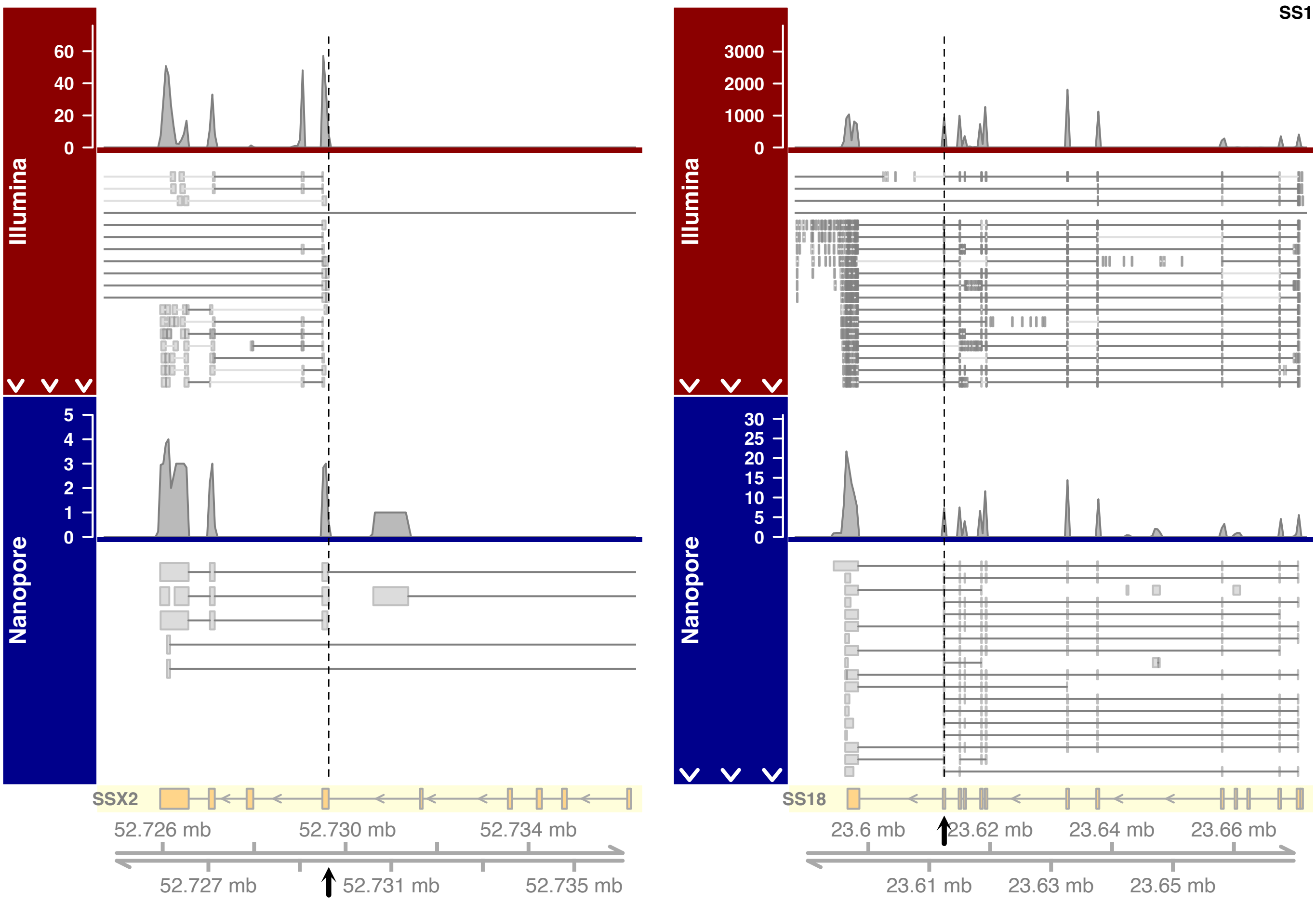


**Fig. S10.** Visualization of short- and long-read alignments to *SSX2* and *SS18* in patient sample SS1. See Fig. S4 for description.
